# The Ca^2+^-Sensitivity of Contraction is Increased in the Left Atrium and Left Ventricle of Patients with Ischemic Heart Failure

**DOI:** 10.64898/2026.06.26.734899

**Authors:** Gregory N. Milburn, Chloe Roth, Jania Bell, Austin Wellette Hunsucker, Marziyeh Pakbaz, Alexandre Lewalle, Steven A. Niederer, Kenneth S. Campbell

## Abstract

**Background:** Ischemic heart failure (IHF) has been shown to impair contractility and disrupt sarcomere function in the left ventricle. Left ventricular failure can cause left atrial dysfunction, which is associated with a greater risk of patient mortality. Despite this, the biochemical and biomechanical characteristics of the left atrium in IHF remain obscure.

**Methods:** Myocardial mechanical properties were measured using permeabilized muscle isolated from the left ventricle (LV) and left atrium (LA) of donors and patients with IHF. Tissue homogenates from these samples were used to measure titin and myosin isoforms as well as the phosphorylation of sarcomeric regulatory proteins. Histology was used to quantify fibrosis in the patients’ left ventricle and left atrium.

**Results:** Length-dependent changes in Ca^2+^-sensitivity were blunted in LV myocardium from patients with IHF. LA myocardium did not show robust length-dependence of Ca^2+^-dependent force. The calcium sensitivity of both LA and LV myocardium was increased in IHF. The maximum force generated by LV but not LA myocardium was decreased in IHF. LA myocardial samples exhibited faster contractile kinetics than LV samples, irrespective of disease. Troponin I phosphorylation decreased in both chambers with IHF.

**Conclusions:** Left atrial IHF myocardium maintained contractile force and displayed increases in calcium sensitivity, which may allow for increased LA contraction under pathological conditions. The increases in calcium sensitivity observed in ischemic myocardium of both chambers are likely driven by decreased phosphorylation of troponin I, which alters thin filament regulation. Conversely, thick filament properties of the left ventricle, such as thick filament protein isoforms and phosphorylation of myosin binding protein-C, displayed chamber-specific differences independent of disease state. These biochemical changes may explain the chamber-specific differences in kinetics and length-dependent properties. Collectively, these biophysical and biochemical data suggest LA remodeling in IHF may assist in increasing LV end-diastolic volume to maintain adequate cardiac output.

## Introduction

Ischemic heart failure (IHF) is the leading cause of death worldwide.^1^ IHF is most often caused by obstructive coronary artery disease, resulting in infarction of the left ventricular (LV) myocardium and subsequent cardiomyocyte death and scar formation. These compositional changes in the LV result in impaired contraction and relaxation, as well as progressive chamber remodeling.^2^ Although the initial ischemic insult is often to the LV, the other chambers of the heart undergo compensatory remodeling in response.^3^ The left atrium (LA) has been shown to dilate and become fibrotic in response to elevated LV filling pressures present during heart failure.^4^ This remodeling of the LA during heart failure disrupts its systolic and diastolic properties, which can hinder atrial function as a pump and conduit, respectively.

As cardiac imaging techniques have advanced, clinical imaging studies have shown that measurements such as LA strain and volume are effective at predicting outcomes in patients with heart failure.^5,6^ These measurements of LA function have been shown to have clinical utility; however, little is known about the changes at the myofilament and tissue levels that may be driving organ-level dysfunction. Prior studies have shown extensive changes in failing LV myocardium, but few studies have examined changes in the mechanical and biochemical properties of LA tissues. Some animal models of heart failure have found that the LA undergoes significant remodeling, including dilation, fibrosis, and impaired function.^7^ While these studies have highlighted the clinical importance and potential mechanisms of LA remodeling in heart failure, the biophysical and biochemical changes in human disease remain largely unexplored. Understanding how myofilament remodeling occurs during disease is important, as sarcomere-specific drugs (myotropes) developed for predominantly LV failure have unintended effects on the LA.^8,9^

To address this gap, we investigated how IHF alters sarcomeric properties in paired human LV and LA myocardium. We found that IHF produced distinct chamber-specific mechanical and biochemical remodeling. Length-dependent activation and maximal force were reduced in IHF LV myocardium, whereas IHF LA myocardium preserved force generation despite increased calcium sensitivity. Decreased troponin I phosphorylation in both chambers of IHF myocardium was associated with increased calcium sensitivity, while differences in thick filament protein properties were chamber-specific. Together, these findings suggest that LA remodeling in IHF may support LV filling and help maintain cardiac output under pathological conditions.

## Methods

### Patient Samples

Myocardial samples were collected from 10 patients with end-stage IHF and 10 age-matched organ donors without a history of cardiac ischemia or cardiovascular disease. Hearts were collected from the operating room and transported on ice to the laboratory. Paired samples from the same patient were taken from the LV and LA and flash frozen in liquid nitrogen. The total time from organ removal to sample freezing was approximately 30 minutes. This method of sample preparation has been shown to preserve many of the contractile and biochemical properties of the myocardium.^10,11^ A detailed description of this biospecimen collection protocol has been published.^12^ All procedures received approval from the University of Kentucky Institutional Review Board (IRB# 46103), and the subjects or their legally authorized representatives provided written informed consent.

### Myocardial Strip Preparation

Multicellular myocardial strips were obtained by mechanical homogenization and chemical permeabilization with 1% v/v Triton before being attached between a motor (model 312B, Aurora Scientific) and a force transducer (model 403, Aurora Scientific), as described.^13^ The sarcomere length of each myocardial strip was adjusted to either 1.94 ± 0.03 or 2.23 ± 0.03 µm in pCa 9.0 solution (pCa= -log_10_[Ca^2+^]). Each strip’s length and cross-sectional area were measured. Force measurements and length changes were normalized to cross-sectional area and muscle length (L_0_), respectively, to account for variations in strip dimensions. Myocardial strip dimensions did not differ between failing and donor groups (Supplemental Figure S1).

### Contractile Function Measurements

To evaluate the force-pCa relationship, myocardial strips were transferred from pCa 9.0 to a pre-activation solution for two minutes and then into activating pCa solutions. Once the strip reached a steady-state force, a series of length changes was applied, after which the strip was returned to pCa 9.0. This sequence was repeated in random order for pCa values ranging from 9.0 to 4.5. All contractile assays were conducted at 37 °C.

Force-pCa curves were generated from each strip by fitting a four-parameter Hill equation:

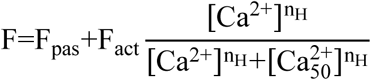

where F_pas_ is the passive force, F_act_ is the maximum active force, n_H_ is the Hill coefficient, and 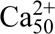 is the concentration of free calcium required to generate half-maximum force. The rate of force recovery, *k*_tr_, was measured by fitting a single exponential function of the form 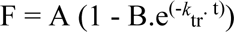 to the force record after a rapid shortening / re-stretch protocol (20% L_0_, 20 ms duration). If a myocardial strip’s maximum force decreased by more than 30% from the beginning to the end of the experiment, it was excluded from analysis.

### Stretch Response Analysis

Additional tissue properties and myosin kinetics were examined by analyzing the transient force response to a step stretch (Supplemental Figure 2). The active component of the force transient was obtained by taking the difference between step stretches (1% L_0_) at minimal (pCa 9.0) and maximal activation (pCa 4.5). Analysis of the force transient in response to a rapid step stretch has been widely discussed in previous publications.^14^ Briefly, the active component exhibits an initial steep increase in force to a peak (P_1_), which then decays to a nadir (P_2_). A single exponential equation, F=A.e^-*k*^_rel_^.t^, was fit to the decay phase where A is the amplitude of P_1_ and *k*_rel_ is the rate at which the force decays to P_2_. Force then redevelops to a new steady state force (P_3_) at a rate *k*_df_, which can be calculated as the linear transformation of the half-time of force redevelopment.^15^

### Immunoblots

Myocardial samples were homogenized using a bead blender (MP FastPrep-24) and then solubilized in a 4 Murea sample buffer. Samples were separated on an 8% SDS-acrylamide gel and then transferred to polyvinylidene difluoride (PVDF) membranes. Membranes were blocked for 60 minutes at room temperature and incubated in primary antibodies overnight at 4 °C. Membranes were incubated in secondary antibodies (1:10000, Invitrogen, 31460) for 60 minutes at room temperature, then washed, and developed in SignalFire ECL (Cell Signaling: 6883S). After imaging, the membranes were stripped and re-probed.

Myosin binding protein-C (MyBP-C) phosphorylation was measured using phosphospecific antibodies for MyBP-C at serines 273, 282, 302 (1:10000, ProSci Inc.), normalized to total MyBP-C (1:5000, Santa Cruz Biotechnology, sc-137237).^16^ Phosphate affinity gels were utilized to measure troponin I (TnI) phosphorylation. This method is similar to traditional immunoblotting but includes the addition of Phos-tag (FujiFilm) to the gel, which allows for the separation of proteins based on both size and phosphorylation status.^17^ Detailed protocols of this method have been described.^18^ These gels were transferred as previously described and probed for TnI (1:5000, HyTest, 4T21). All immunoblots were analyzed using GelBox.^19^

### Titin and Myosin Isoform Gels

Frozen myocardial samples were pulverized in a liquid nitrogen-cooled Dounce homogenizer and solubilized in an 8M urea sample buffer with protease inhibitors to minimize protein degradation. To quantify titin isoforms, protein samples were run on a 1% SDS-agarose gel, as previously described^20^, and stained with Oriole (BioRad: 161-0496) according to the manufacturer’s instructions. Myosin heavy chain (MHC) isoforms were measured by running protein samples on a 7% SDS-PAGE gel at 140 mV for 8 hours.^21^ Gels were then silver-stained (BioRad 1610449) per manufacturer’s instructions. The proportion of titin expressed as the N2BA isoform was calculated as the optical density of the N2BA band divided by the sum of the N2BA and N2B isoform bands. Similarly, the proportion of myosin expressed as the α-MHC isoform was calculated as the optical density of the α-MHC band divided by the sum of the α and β-MHC bands. Band densities were measured using GelBox software.

### Fibrosis Staining and Quantification

Tissue samples were placed into plastic molds filled with optimal cutting temperature (OCT) compound and frozen by submersion in liquid nitrogen-cooled isopentane. Tissue blocks were cryo-sectioned at a thickness of 10 µm in triplicate. Sections from the LV and LA were stained using picrosirius red.^22^

Tissue sections were imaged on a Zeiss Axioscan Z1 slide scanner, and the images were analyzed with a k-means clustering algorithm as previously described.^23^ Relative fibrosis was calculated as the ratio of fibrosis-stained tissue to the total tissue area (myocardium plus fibrosis). Image segmentation and quantification were performed using custom-written functions in MATLAB (MathWorks, Natick, MA, Version R2025a).

### Statistical Analysis

Data were analyzed using linear mixed models in SAS (SAS Institute Inc., Cary, NC, Version 9.4). Linear mixed models included disease and/or region as fixed effects, with the patient’s deidentified code as a random effect. Linear mixed models assumed a compound symmetry covariance structure. The normality of the data was evaluated using Shapiro-Wilk tests in SAS. Pairwise comparisons were made using Tukey-Kramer post-hoc tests. This statistical approach accounts for repeated measurements and paired samples from different regions of the same hearts. Some data are presented in superplots, where the mean measurement for each patient is represented as an opaque symbol, while repeated measures are shown as transparent symbols. Clinical demographics of patients in each group were compared using unpaired t-tests in MATLAB (MathWorks, Natick, MA). Linear regression plots and curve fitting were performed with MATLAB. Continuous variables analyzed via linear regression are presented with slope p-values, 95% confidence intervals, and Pearson’s correlation coefficients (r). P-values less than 0.05 are considered statistically significant. Demographics and echocardiogram measurements are reported as the mean ± standard deviation (SD), while mixed model results are shown as the mean ± standard error of the mean (SEM).

## Results

### Ischemic heart failure blunts length-dependent activation in LV myocardium

Length-dependent activation is described as an increase in maximal force and enhanced calcium sensitivity at longer sarcomere lengths. Compared to donor patients, LV myocardium from patients with IHF did not have a significant increase in maximum active tension or calcium sensitivity with increased sarcomere length (Figure 1). LA myocardium showed no significant length-dependence on calcium sensitivity in both donor and IHF patients (Figure 2). A length-dependent increase in maximum active tension was observed in IHF LA myocardium but not in donor myocardium (Figure 2).

**Figure 1.**
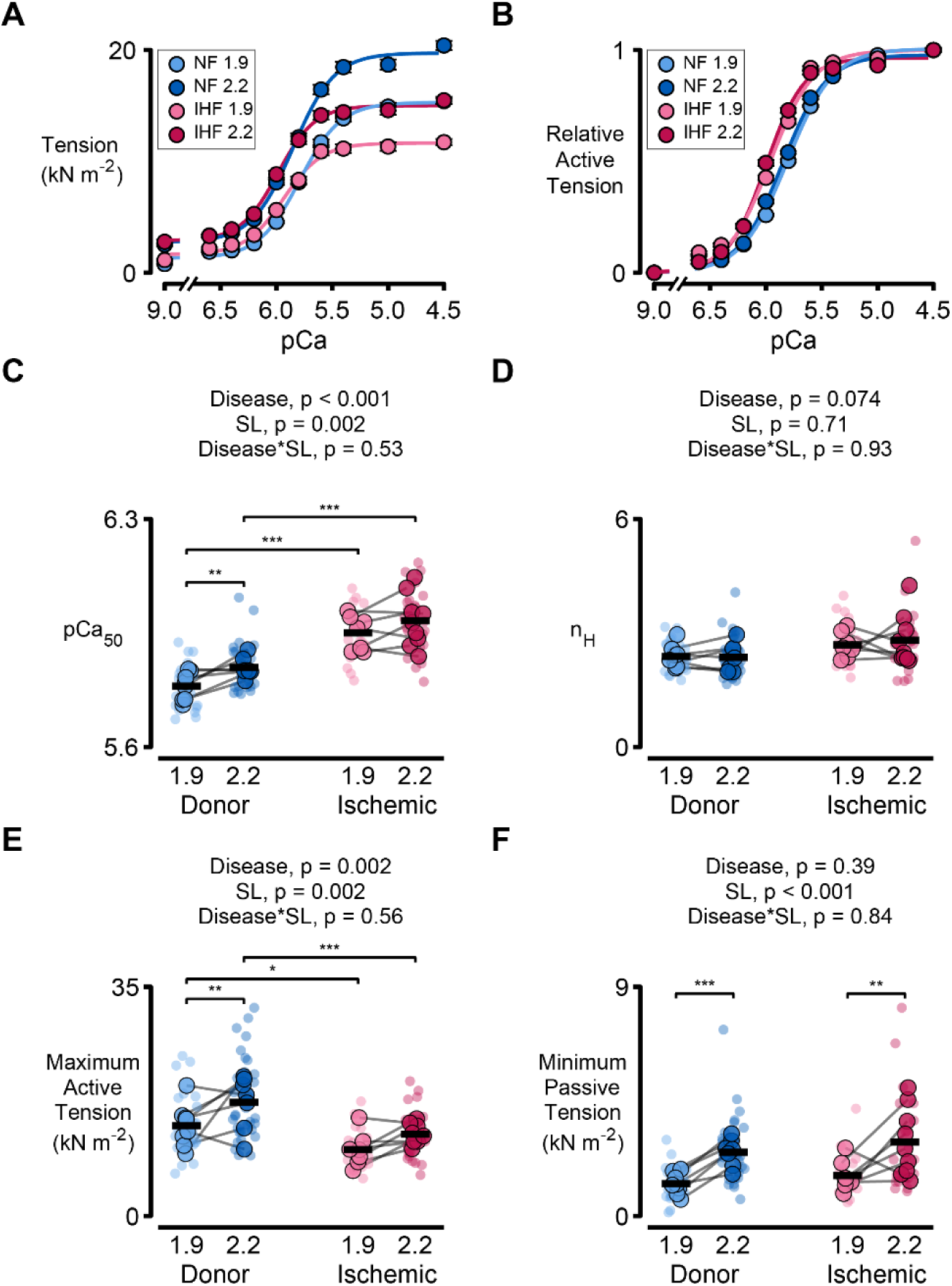
Left ventricle length-dependent activation is blunted in ischemic heart failure. Tension-pCa curves of LV myocardial strips at a sarcomere length of 1.9 and 2.2 μm showing (A) absolute and (B) relative tension as a function of Ca^2+^ concentration (pCa). Superplots show (C) calcium sensitivity (pCa_50_), (D) Hill coefficient (n_H_), (E) maximum active tension, and (F) passive tension. Data were analyzed using linear mixed models. *p < 0.05, **p < 0.01, ***p<0.001

**Figure 2.**
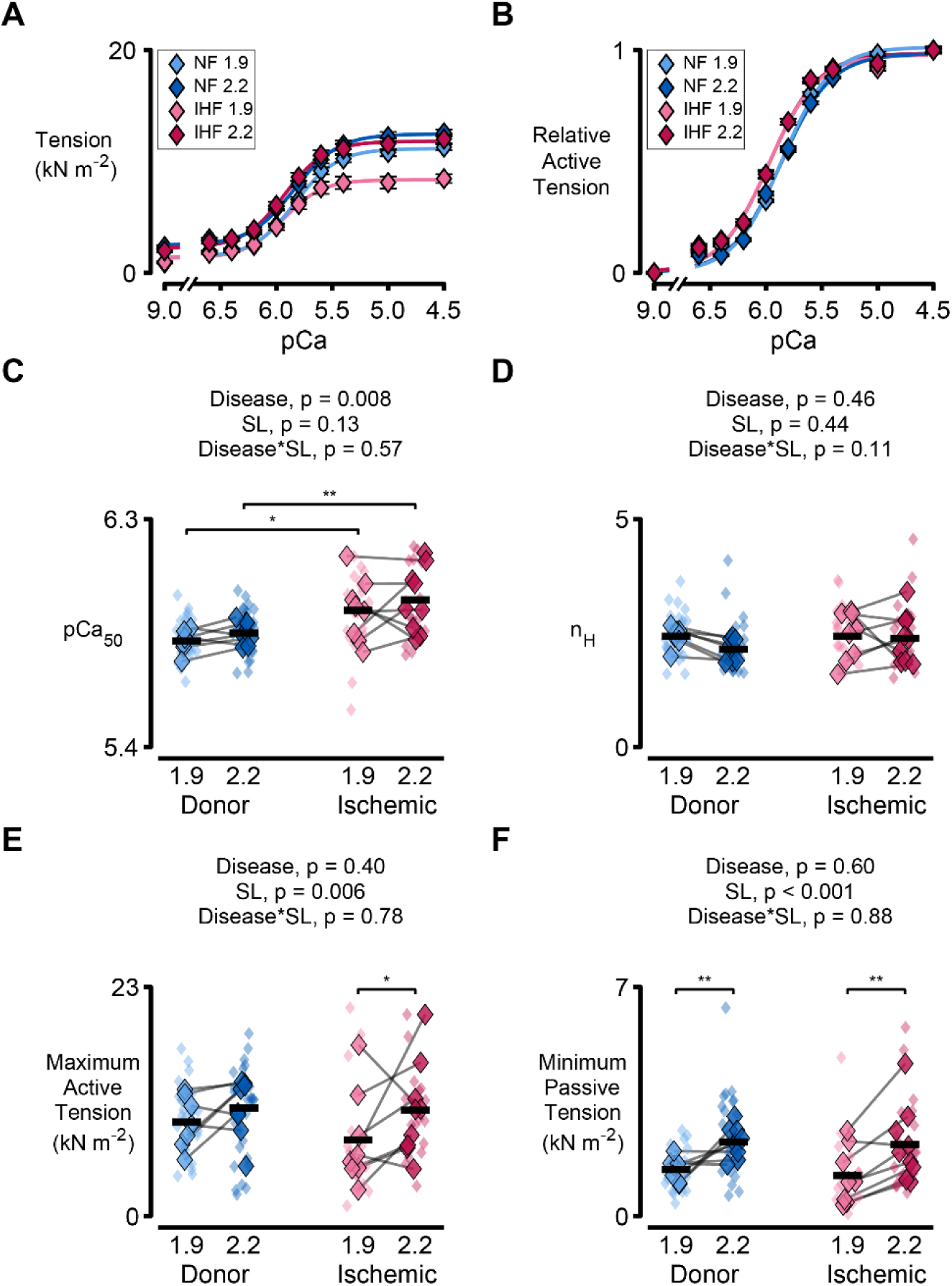
Length-dependent activation is not significant in the left atrium myocardium regardless of disease status. Tension-pCa curves of LA myocardial strips at a sarcomere length of 1.9 and 2.2 μm showing (A) absolute and (B) relative tension as a function of Ca^2+^ concentration (pCa). Superplots show (C) calcium sensitivity (pCa_50_), (D) Hill coefficient (n_H_), (E) maximum active tension, and (F) passive tension. Data were analyzed using linear mixed models. *p < 0.05, **p < 0.01, ***p<0.001

### LV and LA calcium sensitivity is increased in ischemic heart failure

Myocardium from patients with IHF displayed increased calcium sensitivity in both LV and LA tissues, independent of sarcomere length (Figures 1C, 2C). Within the IHF group, calcium sensitivity was significantly increased in LV compared to LA tissues (Figure 3). No difference in calcium sensitivity was observed between the LV and LA of donor myocardium.

**Figure 3.**
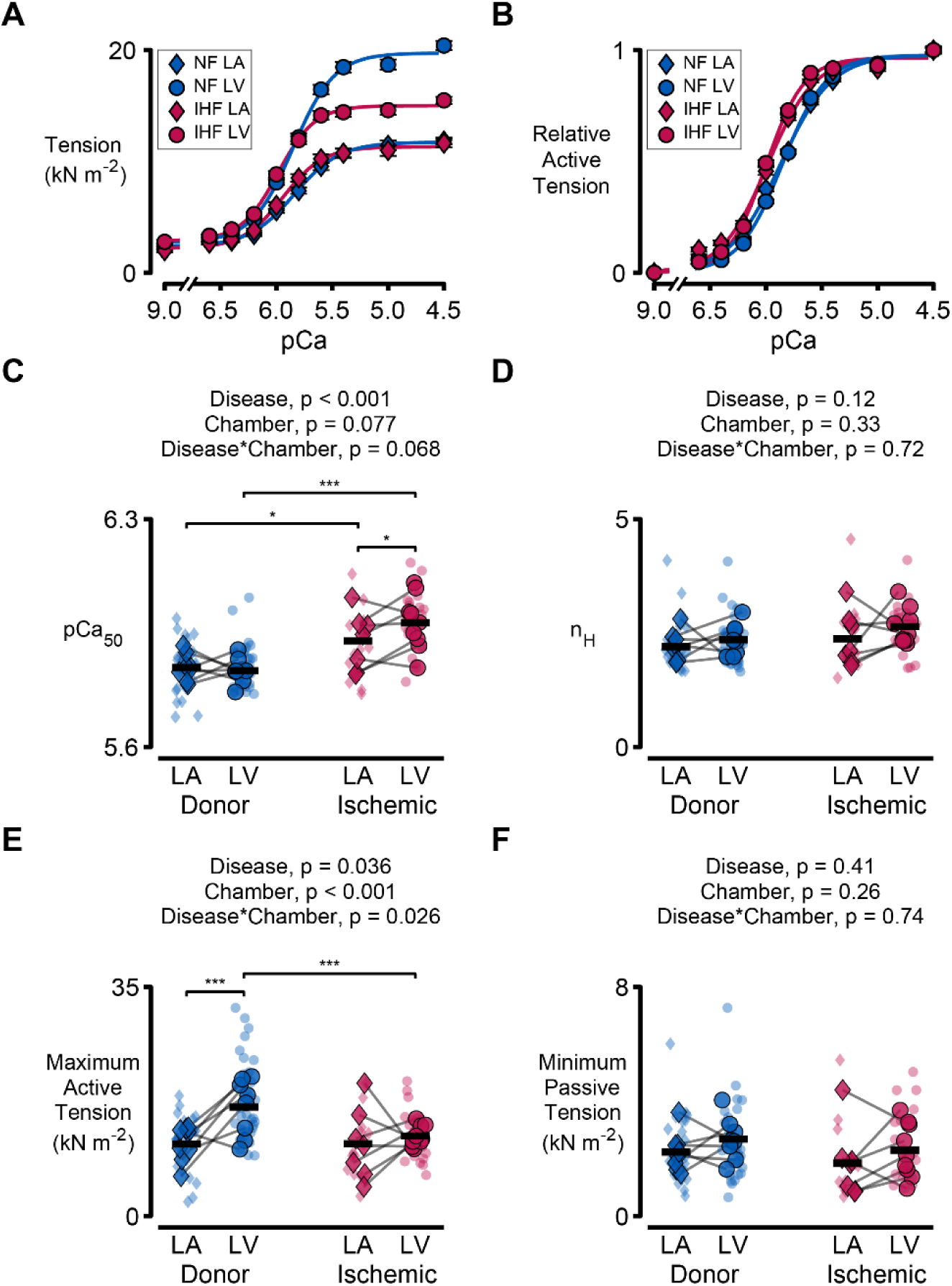
Left atrial force is preserved in ischemic heart failure, while calcium sensitivity is increased in both chambers. Tension-pCa curves of LA and LV myocardial strips at a sarcomere length of 2.2 μm showing (A) absolute and (B) relative tension as a function of Ca^2+^ concentration (pCa). Superplots show (C) calcium sensitivity (pCa_50_), (D) Hill coefficient (n_H_), (E) maximum active tension, and (F) passive tension. Data were analyzed using linear mixed models. *p < 0.05, **p < 0.01, ***p<0.001

### LA force is preserved in ischemic heart failure

There was no significant difference in the maximum active tension generated by LA myocardium between patients with or without IHF (Figure 3E). In contrast, there was a significant decrease in the active tension of LV myocardium from patients with IHF compared to donor myocardium. There was no difference in force production between the LA and LV of failing myocardium.

### Cross-bridge kinetics are faster in the atrium than in the ventricle regardless of disease status

The peak force during the step stretch (P_1_) was increased in both LA and LV tissues from patients with IHF but did not show significant differences between the chambers (Figure 4A). LA preparations displayed faster k*_df_*, k*_rel_*, and k*_tr_* than LV myocardium in both IHF and donor samples (Figure 44B, D, and F). LA tissues also had a larger nadir (P_2_) after step stretches than LV myocardium, regardless of disease status (Figure 4C). No differences in P_3_ were observed between chambers or disease groups (Figure 4E).

**Figure 4.**
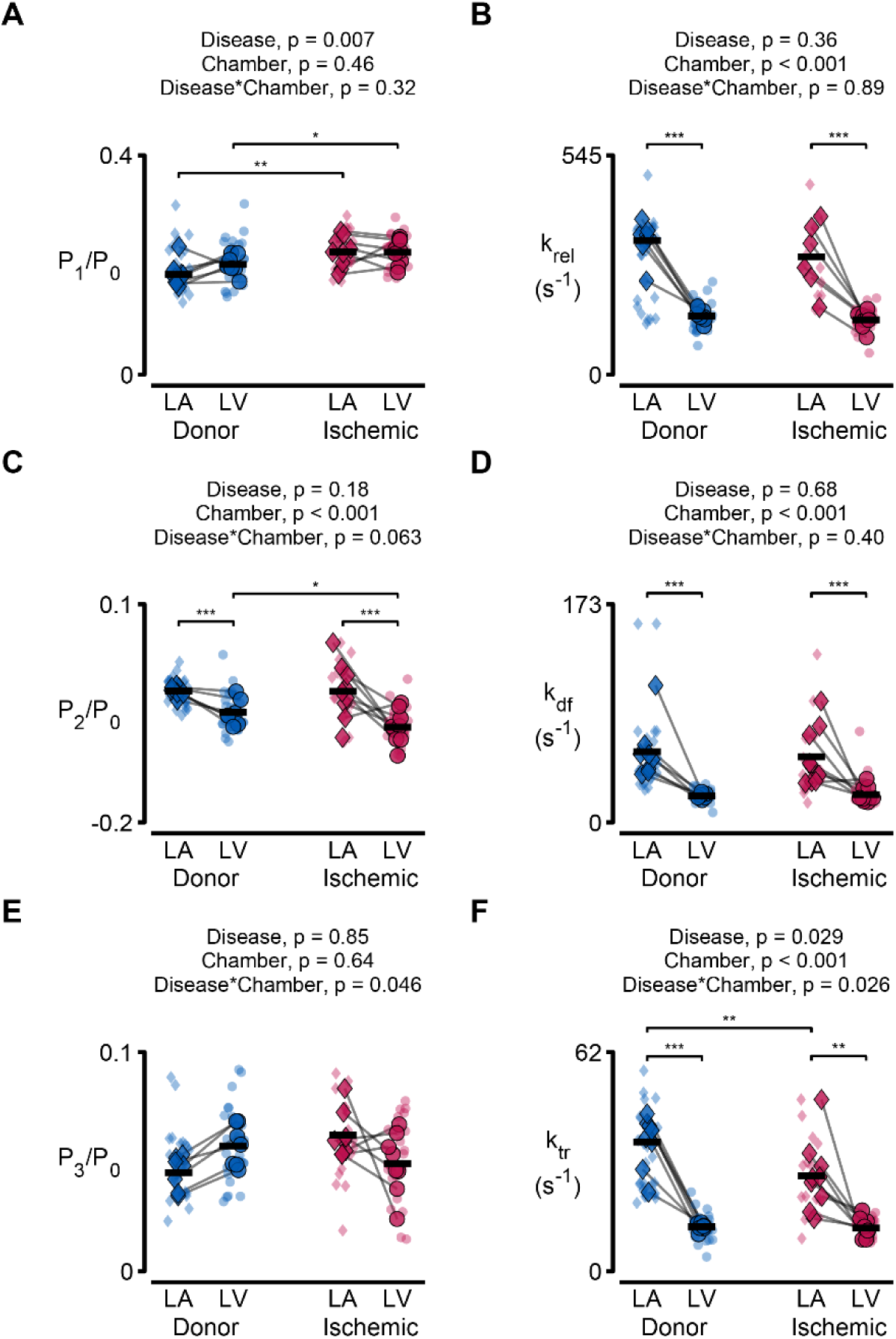
LA myocardium has faster cross-bridge kinetics than the ventricle, regardless of disease status. Parameters calculated from step-stretch force transients or k_tr_ maneuver. Superplots show (A) P_1_, (C) P_2_, and (E) P_3_ normalized to P_0_ and (B) k_rel_, (D) k_df_, (F) k_tr_. Data were analyzed using linear mixed models. *p < 0.05, **p < 0.01, ***p<0.001

### Phosphorylation of MyBP-C and TnI is decreased in ischemic myocardium

Phosphorylation of MyBP-C in the LV was significantly decreased at serines 273, 282, and 302 in IHF myocardium (Figure 5). IHF had no impact on the phosphorylation of the LA MyBP-C at serine 273 and 282, but there was a significant reduction in the phosphorylation of MyBP-C serine 302. TnI phosphorylation decreased in both the ventricle and atrium of ischemic myocardium, with slightly more TnI phosphorylation in donor LV than LA (Figure 6).

**Figure 5.**
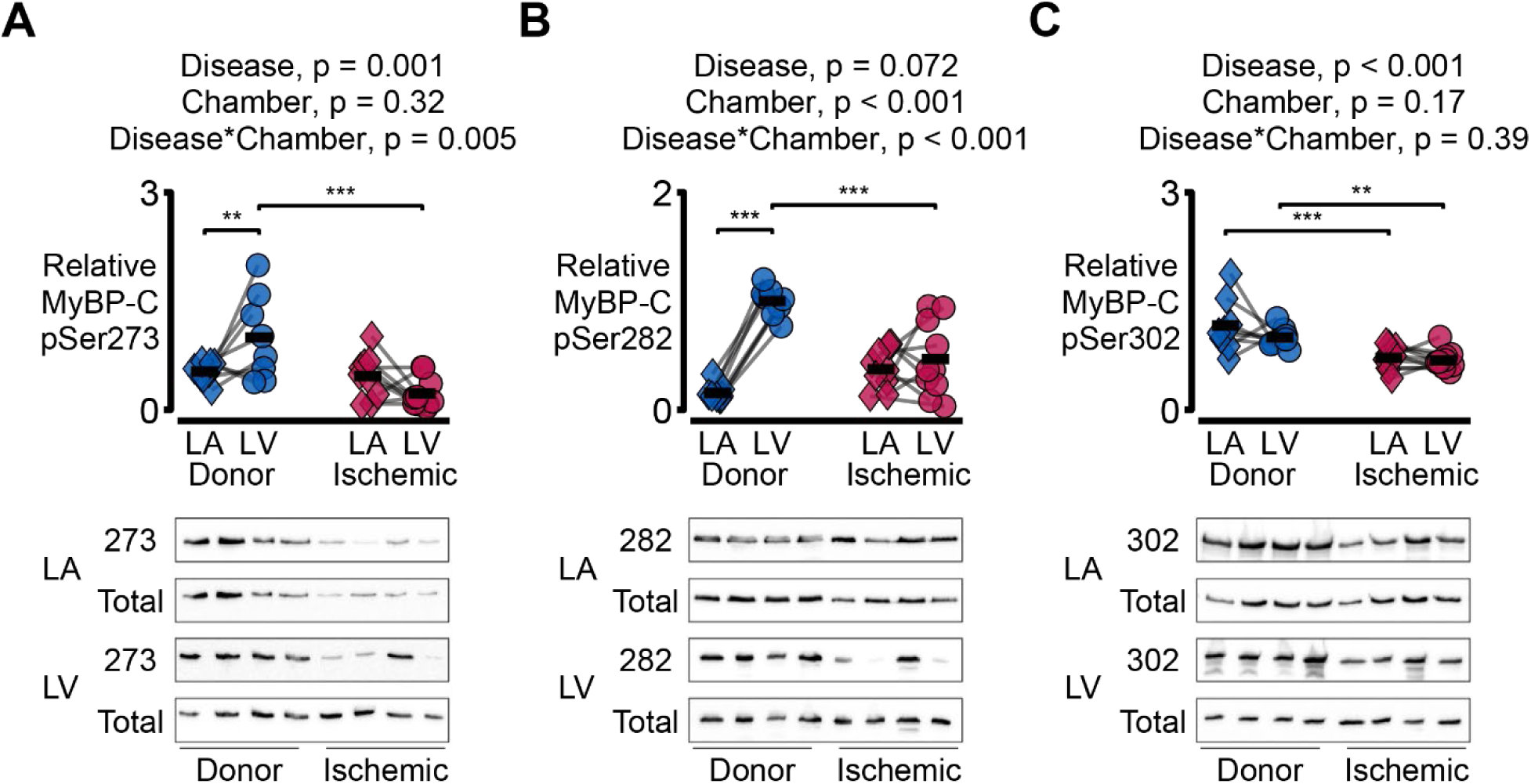
Phosphorylation of Myosin Binding Protein C (MyBP-C) and Troponin I (TnI) is decreased in IHF myocardium. Superplots comparing phosphorylation of MyBP-C at serine (A) 273, (B) 282, (C) 302 relative to total MyBP-C in the LA and LV of IHF and organ donor myocardium. Respective representative immunoblots of total and phosphorylated MyBP-C at serines 273, 282, and 302 in the LA and LV. Values are normalized to the mean relative phosphorylation of donor LV myocardium at each site. Superplot data were analyzed using linear mixed models. *p < 0.05, **p < 0.01, ***p<0.001.

**Figure 6.**
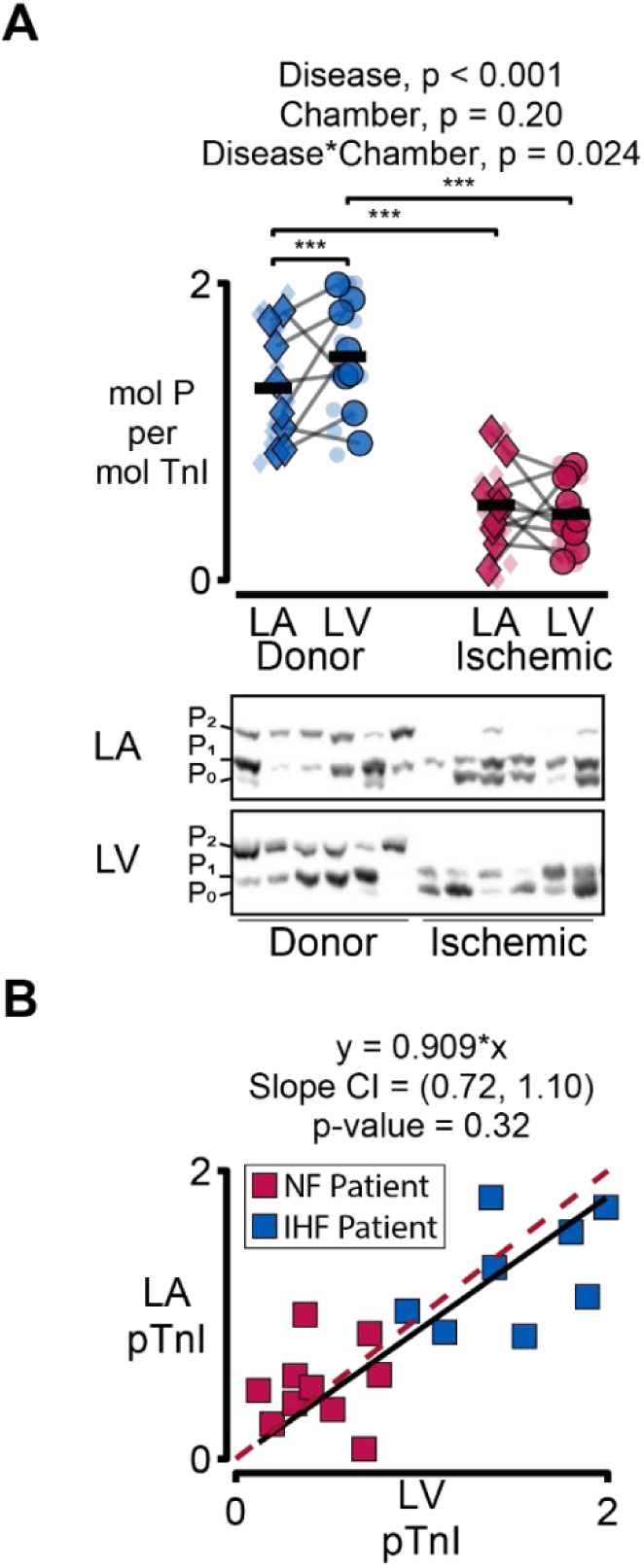
Ischemic heart failure reduces TnI phosphorylation in both the left atrium and ventricle. A) Superplot comparing the amount of phosphorylation in moles of phosphate per mole of TnI in the LA and LV of donor and IHF myocardium. TnI Phos-tag immunoblot showing unphosphorylated (P0), monophosphorylated (P1), and biphosphorylated (P2) TnI. B) Deming regression comparing the phosphorylation of TnI in the LA and LV of IHF and donor patients. The p-value shows the probability of the measured slope differing from 1. The dashed red line represents an identity line where LV TnI equals LA TnI. Superplot data were analyzed using linear mixed models. *p < 0.05, **p < 0.01, ***p<0.001.

### Fibrosis, α-myosin, and N2BA titin are increased in LA myocardium

The percentage of α-myosin increased in LA compared to LV myocardium in both IHF and donor myocardium (Figure 5A). The proportion of titin expressed as the N2BA isoform was increased in the atria compared to the ventricle in both ischemic and donor myocardium (Figure 7B). In myocardium from donor patients, LA samples showed significantly greater fibrosis than their respective LV (Figure 7C,D). The LV of IHF samples showed increased fibrosis compared with donor samples; however, there was no difference in LA fibrosis between patients and donors (Figure 7D).

**Figure 7.**
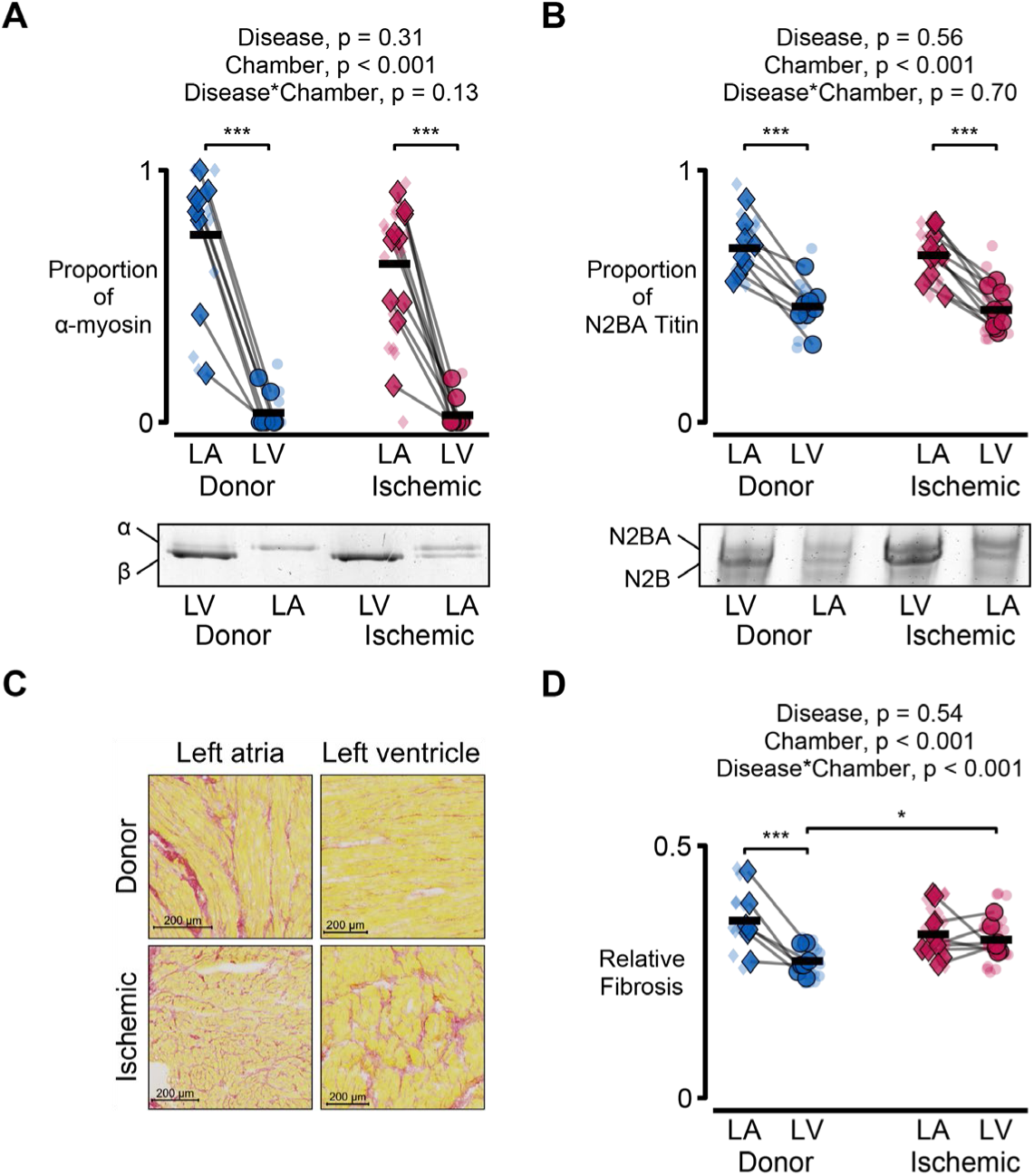
Fibrosis, α-MHC, and N2BA titin are increased in LA myocardium. (A) Plot comparing the proportion of α myosin of the LA and LV in IHF and donor myocardium. (B) Plot comparing the proportion of LA or LV N2BA titin in IHF and donor myocardium. (C) MHC gel showing myosin isoform content in the LA or LV of IHF and donor myocardium. (D) Representative titin gel of the LA and LV of IHF and donor myocardium. (E) Representative image of picrosirius red staining of the left heart in IHF and donor samples. (F) Superplot comparing LA and LV fibrosis content in IHF and donor hearts. Data were analyzed utilizing linear mixed models. *p < 0.05, **p < 0.01, ***p<0.001.

**Figure 8.**
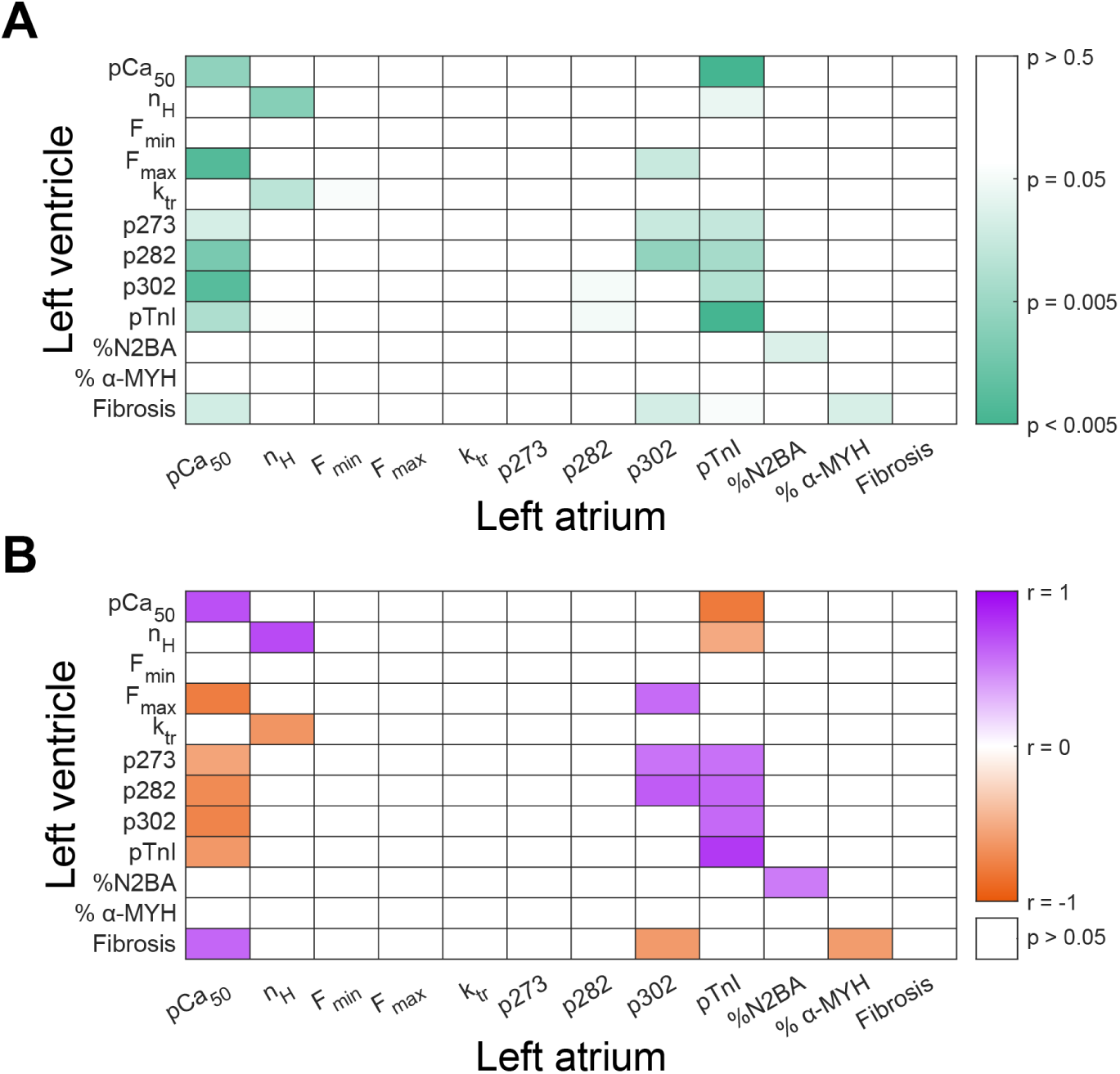
TnI phosphorylation is tightly coupled between the left atrium and ventricle. (A) Heatmap showing p-values of linear regressions between paired mechanical and biochemical assays on the left ventricle and atrium. (B) Heatmap showing correlation coefficients (r) from linear regressions shown in panel A.

## Discussion

In this study, we showed that IHF results in chamber-specific alterations to mechanical and biochemical properties of LA and LV myocardium. While IHF increases calcium sensitivity in both LA and LV, force was reduced only in LV myocardium. These changes in muscle mechanics appear to be driven by post-translational modifications of sarcomeric proteins and extracellular matrix remodeling. To our knowledge, this is the first study to examine how the mechanical and biochemical properties of the sarcomere are impacted by ischemic heart failure in paired LA and LV samples.

### Patient Characteristics

The donor cohort was selected from organ donors not viable for donation but with a non-cardiac cause of brain death and no history of cardiovascular disease. The patients with IHF were collected from patients receiving heart transplantation. Efforts were made to minimize the confounding effects of age, sex, BMI, and diabetic status between the failing and donor cohorts. There was no significant difference in age, BMI, or HbA1C between the patient cohorts (Supplemental Table S1). Consistent with heart failure remodeling, there was a significant reduction in ejection fraction and an increase in LV internal diameter in the ischemic patient cohort. While some patients in the ischemic cohort had multivessel disease, all patients had pathologic involvement of the left anterior descending artery (Supplemental Table S2).

### Length-dependent activation is blunted in ischemic LV myocardium

The Frank-Starling mechanism is a fundamental property of the heart by which increased end-diastolic volume leads to increased contractility.^24^ At the sarcomeric level, length-dependent activation (LDA) links increased sarcomere length to increased force and calcium sensitivity.^25^ We observed a reduction of LDA in ischemic LV myocardium; neither maximal force per cross-sectional area nor Ca^2+^-activated force increased with sarcomere lengthening (Figure 1E and 1C). This reduction in length-dependent contractility in patients with IHF may compromise synchronicity of contraction and the heart’s ability to adjust to changes in preload.^26^ Taken together, these changes may contribute to exercise intolerance, a hallmark feature of advanced heart failure.

While the full mechanism of LDA continues to be explored, the post-translational status of sarcomeric proteins is known to be an important factor. Phosphorylation of Myosin Binding Protein-C (MyBP-C), a thick filament-associated regulatory protein, modulates cross-bridge interaction by amplifying length-dependent increases in Ca^2+^ sensitivity.^16^ We have previously shown that MyBP-C phosphorylation is reduced in patients with heart failure, independent of etiology.^23,27^ Consistent with these findings, LV myocardium from patients with IHF exhibited significant reductions in MyBP-C phosphorylation at all Protein Kinase A (PKA)-targeted sites—serines 273, 282, and 302 (Figure 5A, 5B, and 5C). These alterations in thick filament regulation in the failing LV likely contribute to the impaired LDA observed in IHF, thereby limiting the ischemic heart’s functional reserve.

### LA myocardium displays reduced length-dependent properties

Current evidence for LDA shows that both thin and thick filament activation is enhanced at longer sarcomere lengths. Filament activation is modulated by regulatory properties such as sarcomeric protein phosphorylation and isoform expression.^28,29^ These properties have been widely explored in heart failure and implicated in dampening LDA in the failing LV; however, their status in the LA is less well characterized.

We found that LA myocardium displayed minimal LDA in both IHF and donor tissue (Figure 2E). A modest but significant increase in maximum active force was observed with increased sarcomere length in IHF LA myocardium, but calcium sensitivity did not change with sarcomere length (Figure 2C). This was unexpected, as both length-dependent contractility and calcium sensitivity were reduced in failing LV myocardium but maintained in LV myocardium of non-failing hearts. Thus, robust LDA appears to be largely absent in the human LA regardless of disease state.

Blunted LDA in the LA compared to the LV may be attributed to chamber-specific differences in thick filament properties, including isoform composition and post-translational modifications. In patients with IHF, LA myocardium maintained MyBP-C phosphorylation at serines 273 and 282 (Figure 5), in contrast to the IHF LV, which displayed low MyBP-C phosphorylation at all PKA-phosphorylated sites (Figure 5). Because LDA is reduced in the LA regardless of disease state, length-dependent properties may not be solely disease-driven but rather influenced by the cardiac chamber-specific differences in sarcomeric proteins.

On the sarcomeric level, the healthy atria and ventricles exhibit differences in thick filament protein isoforms which may contribute to the discrepancy in LDA between the LV and LA. Titin, a large sarcomeric protein that acts as a molecular spring, possesses a mechanosensing mechanism whereby increased strain can influence myosin head availability for thin filament binding.^30,31^ This mechanosensing mechanism can be affected by titin’s stiffness, which is dependent on its isoform expression.^32^ Titin isoform expression is a primary determinant of myocardial stiffness because its different isoforms, N2B and N2BA, differ in extensibility and passive tension generation^33^. LA myocardium had a greater proportion of the more compliant titin isoform, N2BA, compared to LV tissue (Figure 7B, D). This increased titin compliance may reduce force-dependent recruitment of myosin heads at longer sarcomere lengths, thereby attenuating LDA.^32^ Likewise, myosin heavy chain (MHC) isoform expression also exhibits chamber-specific properties. LA myocardium displayed a substantially higher proportion of α-MHC than the LV, which was composed predominantly of β-MHC (Figure C, A). These differences in myosin and other thick filament proteins within the atria produce distinct mechanical and regulatory properties, including a decreased proportion of myosin heads in the super-relaxed state.^34^ Differences in these mechanisms of thick filament regulation may contribute to the divergent LDA properties measured between chambers.

The reduction of LDA observed in the atrium aligns strongly with its physiological role. In individuals with normal diastolic function, approximately 40% of total LV filling volume occurs during the LA reservoir phase, 35% during passive conduit flow, and only 25% during active LA contraction.^35^ Thus, LDA may play a smaller role in LA function compared to the LV, as the atrium does not rely heavily on preload-dependent contractile augmentation. Additionally, the LA-specific protein myosin binding protein H-like (MyBP-HL) may contribute to reduced LDA. MyBP-HL has been implicated in modulating LA relaxation and may further attenuate LDA by reducing the availability or functional impact of MyBP-C phosphorylation on thick filament activation.^36^ Overall, LDA is blunted in LA myocardium compared to LV tissue regardless of disease state, potentially reflecting chamber-specific differences in thick filament properties. Specifically, the LA’s increased titin compliance, α-MHC predominance, and LA-specific regulatory proteins may all contribute.

### Ischemic LA myocardium has increased calcium sensitivity with preserved force production

The role of calcium in the development of active force is fundamental to cardiac contraction. During systole, binding of calcium to Troponin C (TnC) on the troponin complex induces a conformational change of tropomyosin, exposing myosin binding sites on actin and permitting cross-bridge cycling. The relationship between calcium concentration and force production is classically characterized by the force-pCa curve, in which steady-state force is plotted against the negative logarithm of the calcium concentration (pCa = −log[Ca²⁺]). A leftward shift of this curve indicates increased myofilament calcium sensitivity (i.e., a higher pCa₅₀, the calcium concentration at which half-maximal force is achieved), meaning that less calcium is required to generate a given level of force.

In the present study, both IHF LV and LA myocardium demonstrated increased calcium sensitivity, as evidenced by a leftward shift of the force-pCa relationship (Figure 3A and 3B). It has been shown that failing myocardium exhibits an increase in calcium sensitivity as a possible compensatory response to restore active force.^37^ Interestingly, IHF LA myocardium exhibited increased calcium sensitivity without a concomitant reduction in maximal force production, which was observed in IHF LV myocardium. This combination suggests that at physiological calcium concentrations, IHF LA myocardium would be expected to generate greater active force than non-IHF LA tissue, as the leftward shift of the force-pCa curve places the operating point of the myofilaments at a higher relative force for any given submaximal calcium level.

Phosphorylation of troponin I (TnI) at Ser22/23 by protein kinase A (PKA) reduces the calcium sensitivity of troponin C, playing a critical role in tuning proper myocardial relaxation. In the failing human heart, TnI phosphorylation levels are markedly reduced, and this dephosphorylation has been consistently associated with increased myofilament calcium sensitivity. In the present study, both IHF LA and LV myocardium exhibited reduced TnI phosphorylation, which could explain the observed increase in calcium sensitivity (Figure 6).

In heart failure, increased myofilament calcium sensitivity has been proposed as a compensatory response to augment force production in the setting of depressed contractility and impaired calcium handling.^38^ Interestingly, while IHF LV myocardium in the present study showed a reduction in maximal force, the IHF LA did not. This is likely due to differences in fibrotic remodeling between the two chambers; IHF LV myocardium exhibited a significant increase in fibrosis, whereas no difference was observed between the LA groups. Following myocardial ischemia, cardiomyocyte death triggers a reparative fibrotic response in which activated myofibroblasts deposit collagen to maintain structural integrity and prevent ventricular rupture.^39^ Although initially protective, this replacement of contractile cardiomyocytes with non-contractile fibrotic tissue results in less force generation per unit volume in multicellular preparations. The absence of significant fibrosis in the IHF LA may therefore explain why force production was preserved in this chamber despite the ischemic insult, while the substantial fibrotic burden in the IHF LV contributed to its reduced force output.

### Cross-bridge kinetics are largely unchanged in disease

Broadly, cross-bridge kinetics refers to the speed of the actin-myosin interactions within myofilaments that allow the heart to produce the force required to pump blood. In these experiments, crossbridge kinetics were relatively independent of disease status; however, the cardiac chamber had a large effect (Figure 4B, D, and F). LA kinetics were faster compared to the LV kinetics in both IHF and donor myocardium. This is likely driven by an increase in α-MHC in the LA (Figure 4A), which has a faster rate of ATP hydrolysis than β-MHC, and is predominantly expressed in the ventricle.^40^ Additionally, IHF atria have a slightly faster k_tr,_ which could be attributed to faster ATP turnover from α-MHC overall (Figures 7A, F). This may also reflect subtle changes in post-translational modifications of myosin itself, including phosphorylation or acetylation, which have been reported to affect cross-bridge kinetics.

### Increased compliant N2BA titin in the LA may accommodate increased LV end-diastolic pressures

Elevated LV end-diastolic pressure in IHF progressively impairs LA reservoir function by exposing the LA to chronic pressure overload. ^41^ This elevated pressure can be transmitted into the pulmonary vasculature, resulting in pulmonary edema and symptoms of congestive heart failure. LA compliance accommodates increased filling pressures, thereby protecting the pulmonary circulation.^42^

We observed greater expression of the more compliant isoform of titin, N2BA, in the LA compared to the LV in both IHF and donor myocardium (Figure 7B). This is consistent with prior reports demonstrating predominant N2BA expression in mammalian atria.^43^ In the setting of IHF, increased N2BA expression may enhance LA compliance and reservoir capacity, allowing the LA to reduce transmission of elevated pressures into the pulmonary circulation.

### Fibrosis is increased in the ventricle but not the atria in ischemic heart failure

In a healthy heart, the atria contain a higher relative density of fibroblasts, rendering it more susceptible to profibrotic signaling.^44^ Consistent with this, donor patients had significantly greater fibrosis in the LA than in the LV (Figure 7D). The LA operates at substantially lower pressures than the LV but possesses a markedly thinner wall. This baseline LA fibrosis may play a structural role in maintaining transmural chamber integrity when the myocardium is subjected to high degrees of stretch. When considered alongside the titin data, this suggests a mechanical phenotype: the LA, with greater expression of the compliant N2BA titin isoform, may remain relatively compliant over physiological sarcomere lengths, facilitating reservoir and conduit function. However, at longer sarcomere lengths or higher filling pressures, increased fibrosis dominates passive stiffness, resulting in a steeper rise in passive tension.^45^ This dual behavior, compliant within the physiological operating range but stiffer at pathological stretch, may help the LA accommodate routine volume fluctuations while limiting excessive dilation during chronic pressure overload.

### LA and LV have similar thin filament properties but divergent thick filaments

Phosphorylation and isoform composition of sarcomeric proteins can enhance length-dependent activation and tune overall cardiac performance. The interaction between sarcomeric thin and thick filaments is essential for myocardial function in both the atria and ventricles. Coordination of these filaments is regulated in part through phosphorylation of the regulatory proteins MyBP-C and TnI. Previous studies have shown that MyBP-C and TnI phosphorylation are decreased in the LV of failing hearts, which may contribute to reduced cardiac function.^46^ Consistent with these findings, we observed that LV MyBP-C phosphorylation at serines 273, 282, and 302 and overall TnI phosphorylation were significantly lower in IHF hearts (Figures 5A).

In contrast, the LA displayed more selective changes in MyBP-C phosphorylation compared to the LV. Only phosphorylation at serine 302 was low in IHF LA myocardium, whereas MyBP-C phosphorylation was decreased at all tested sites in IHF LV myocardium. PKA-mediated phosphorylation of MyBP-C at serines 273, 282, and 302 accelerates cardiac contractility; however, serine 302 can also be phosphorylated by other kinases, most notably Protein Kinase C (PKC).^16^ This suggests that different kinases may differentially target MyBP-C in the LA compared to the ventricle. Additionally, chamber-specific differences in phosphatase (PP1 and PP2A) regulation have been shown to contribute to distinct disease phenotypes and even exhibit phospho-site specificity for MyBP-C. ^47^ These findings may suggest that these signaling pathways are differentially affected by ischemic disease in the LA compared to the LV.

LA TnI phosphorylation was also reduced in IHF hearts, mirroring the changes observed in LV TnI (Figure 6B). Both TnI and MyBP-C serine 302 phosphorylation have been shown to regulate thin filament activation.^48^ These data suggest that ischemia disrupts thin filament regulation in both the LA and LV, while thick filament regulation is robustly impacted only in the LV.

Titin and MHC isoform analysis further supports divergent thick filament properties between chambers independent of disease. Across both donor and IHF hearts, LA myocardium exhibited higher α-MHC and N2BA titin expression relative to the LV (Figure 7A, B). These chamber- and disease-specific phosphorylation and isoform patterns are summarized in the heat maps shown in Figure 6, which illustrate shared reductions in thin filament phosphorylation across chambers alongside pronounced chamber-specific differences in thick filament composition.

Together, these findings indicate that thin filament regulatory changes are disease-driven and coupled between chambers, whereas thick filament properties appear to be more heavily influenced in a chamber-specific manner. This suggests the LA and LV share biochemical responses to ischemic disease at the thin filament level but diverge in thick filament composition and regulation.

### Limitations

Our lab has been biobanking cardiac samples for over 15 years. Of the patients with ischemic heart failure in our biobank, only 10% are female. This is relatively congruent with other population-based studies.^49,50^ This study included myocardium from patients with IHF, all of whom were male. Although these demographics are representative of our medical center’s patient population with IHF, access to female ischemic myocardium could elucidate sex-dependent mechanisms or differences.

While all patients with IHF included in this study had involvement of the LAD, the spatial relationship to any prior ischemic insult and the myocardial sample used is not well characterized. Additionally, the study had limited access to LA tissue from patients with atrial fibrillation or arrhythmias. Future studies should be conducted to examine how proximity to ischemic damage and arrhythmia may impact sarcomeric mechanical or biochemical properties.

## Conclusions

This study sought to define disease-driven changes to the left atrial and ventricular myocardium in patients with ischemic heart failure. The most prominent finding was that thin filament regulation is altered in disease, while the thick filament modulation is chamber-dependent. Additionally, IHF hearts have blunted length-dependent activation in the ventricle, but not in the atria. This is likely a result of reduced MyBP-C phosphorylation in the left ventricle. Both chambers in IHF hearts displayed increases in calcium sensitivity, which may be attributed to reduced phosphorylation of TnI.

## Acknowledgements

The authors would like to thank the patients and their families for donating cardiac samples to the Gill Cardiovascular Biorepository, which has supported this research.

## Supplemental Material

Tables S1, S2

Figures S1-S4

## Abbreviations

pCa_50_: calcium sensitivity
n_H_: Hill coefficient
LDA: Length-dependent activation
TnI: Troponin I
MyBP-C: Myosin binding protein-C
MHC: Myosin heavy chain

