## Supplemental Materials for "The Ca^2+^-Sensitivity of Contraction is Increased in the Left Atrium and Left Ventricle of Patients with Ischemic Heart Failure"

### Contents:


| Supplemental Material | Page |
|---|---|
| Supplemental Table 1. Summary Group Characteristics | **2** |
| Supplemental Table 2. Individual Patient Characteristics | **3** |
| Supplemental Figure 1. Myocardial strip dimensions did not differ between ischemic and non-failing samples | **4** |
| Supplementary Figure 2. Left ventricle and atria have different force transients in response to step stretches | **5** |
| Supplementary Figure 3. Left ventricle cross-bridge kinetics are slower at longer sarcomere length | **6** |
| Supplementary Figure 4. Sarcomere length and disease status do not significantly alter left atrial cross-bridge kinetics | **7** |

Supplemental Table 1. Summary Group Characteristics

| | Donor (n = 10) | Ischemic HF (n = 10) | p-value |
|---|---|---|---|
| Age | 52.3 ± 11.8 | 62.4 ± 9.8 | 0.053 |
| BMI | 26.5 ± 3.8 | 27.3 ± 3.7 | 0.63 |
| HbA1C | 5.4 ± 0.6 | 6.5 ± 2 | 0.14 |
| LVEF (%) | 55.4 ± 12.4 | 22.3 ± 5.1 | < 0.001 |
| LVIDd (cm) | 4.5 ± 0.5 | 6.2 ± 0.8 | < 0.001 |
| LVPWd (cm) | 1.1 ± 0.2 | 0.9 ± 0.1 | 0.064 |

Patient clinical characteristics and demographics. Values are reported as mean ± standard deviation. BMI = body mass index; HbA1C = hemoglobin A1C; LVEF = left ventricle ejection fraction; LVIDd = left ventricle interior diameters at diastole; LVPWd = left ventricle posterior wall thickness; HF = heart failure.

Supplemental Table 2. Individual Patient Characteristics

| Patient ID | Patient Group | Race | Sex | Age | Donor cause of death | Diabetic | BMI | HbA1C | LVIDd (cm) | LVPWd (cm) | Ejection fraction | Coronary vessel(s) with disease |
|---|---|---|---|---|---|---|---|---|---|---|---|---|
| 2508D | Donor | White | Male | 41 | CVA/Stroke | No | 32.1 | 5.3 | 5.2 | 1.4 | 31 | - |
| 2B487 | Donor | White | Male | 42 | TBI | No | 31.0 | 5.3 | 4.1 | 0.99 | 55 | - |
| 2DDC0 | Donor | White | Male | 49 | TBI | No | 24.4 | 5.2 | 5 | 0.8 | 56 | - |
| 30B2B | Donor | White | Male | 71 | Anoxic brain injury | No | 24.2 | 5.2 | 4 | 1.2 | 66 | - |
| 5245C | Donor | White | Male | 55 | CVA/Stroke | No | 19.7 | 4.8 | | | | - |
| B23E3 | Donor | White | Male | 50 | CVA/Stroke | No | 29.4 | 5.5 | 4.2 | 0.99 | 55 | - |
| BC90C | Donor | White | Female | 39 | Anoxic brain injury | No | 24.1 | 5.2 | 4.4 | 0.98 | 55 | - |
| BE318 | Donor | White | Male | 46 | CVA/Stroke | Yes | 24.5 | 7.0 | 4.7 | 1.1 | 70 | - |
| 8CB30 | Donor | White | Female | 61 | TBI | No | 28.7 | - | - | - | - | - |
| A9200 | Donor | Black or African American | Male | 70 | CVA/Stroke | Yes | 26.5 | 5.4 | - | - | - | - |
| 0DC6E | Ischemic | Black or African American | Male | 55 | - | Yes | 32.9 | 6.1 | 4.9 | 1.1 | 32.9 | LAD, RCA, PDA |
| 2256F | Ischemic | Black or African American | Male | 64.26 | - | No | 25.7 | 5.7 | 6.7 | 1.1 | 14.8 | LAD, PDA |
| 2D65C | Ischemic | White | Male | 63 | - | Yes | 25.6 | 8.5 | 6.3 | 0.78 | 21.9 | LAD,RCA, Cx |
| 5845F | Ischemic | White | Male | 71 | - | No | 30.6 | 6.0 | 6.7 | 1 | 20 | LAD |
| 9664E | Ischemic | White | Male | 42 | - | No | 27.4 | 5.5 | 5.8 | 0.75 | 21 | LAD,RCA |
| AF1FF | Ischemic | White | Male | 75 | - | No | 32.8 | 5.2 | 7.1 | 0.83 | 23 | LAD |
| D6A67 | Ischemic | White | Male | 53 | - | No | 23.0 | 5.6 | 6.4 | 0.77 | 20 | LAD,RCA, Cx |
| DFEFA | Ischemic | White | Male | 67 | - | No | 22.7 | 5.3 | - | - | - | LAD,RCA, Cx |
| FAC13 | Ischemic | White | Male | 65 | - | No | 25.5 | 5.8 | 7.1 | 0.79 | 20 | LAD |
| FCFAD | Ischemic | White | Male | 69 | - | Yes | 26.5 | 11.7 | 5.2 | 0.96 | 27 | LAD,RCA |

CVA = cerebrovascular attach; TBI = traumatic brain injury; BMI = Body mass index; HbA1C = hemoglobin A1C; LVIDd = left ventricle interior diameter at diastole; LVPWd = left ventricle posterior wall thickness; LAD = left anterior descending; RCA = right coronary artery; PDA = posterior descending artery; Cx = circumflex

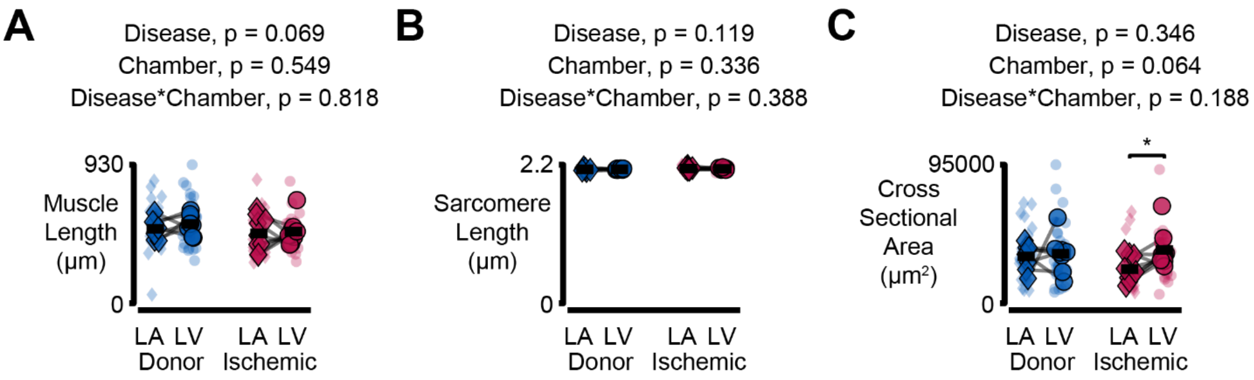


Supplemental Figure 1. Myocardial strip dimensions did not differ between ischemic and non-failing samples

Superplots comparing multicellular myocardial strip (A) length, (B) sarcomere length, and (C) cross-sectional area between ischemic heart failure and non-failing samples at a sarcomere length of 2.2 microns. Transparent symbols represent individual muscle preparations, while opaque symbols represent the mean of each patient. Data were analyzed using linear mixed models. $^*p < 0.05$, $^{**}p < 0.01$, $^{***}p<0.001$

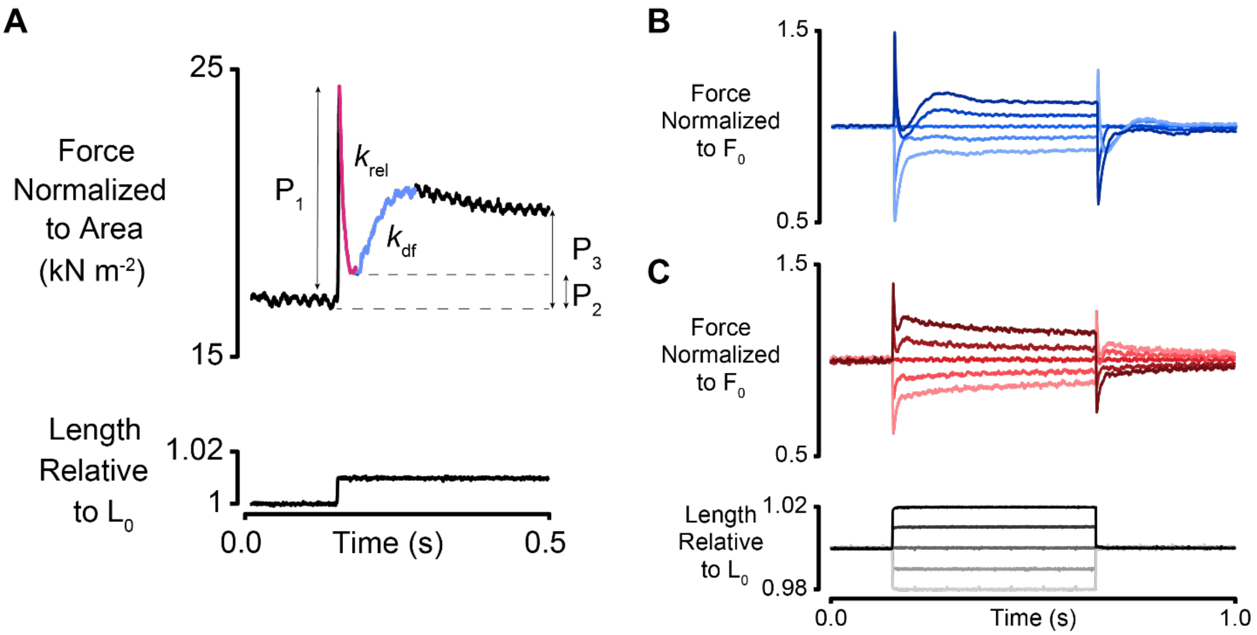


Supplementary Figure 2. Left ventricle and atria have different force transients in response to step stretches

To examine cross-bridge dynamics, (A) preparations at steady state force were rapidly lengthened 1% of the initial muscle length ($L_0$) and the resulting force transient recorded. The force transient in response to step length changes differs between (B) left ventricle and (C) left atria myocardium.

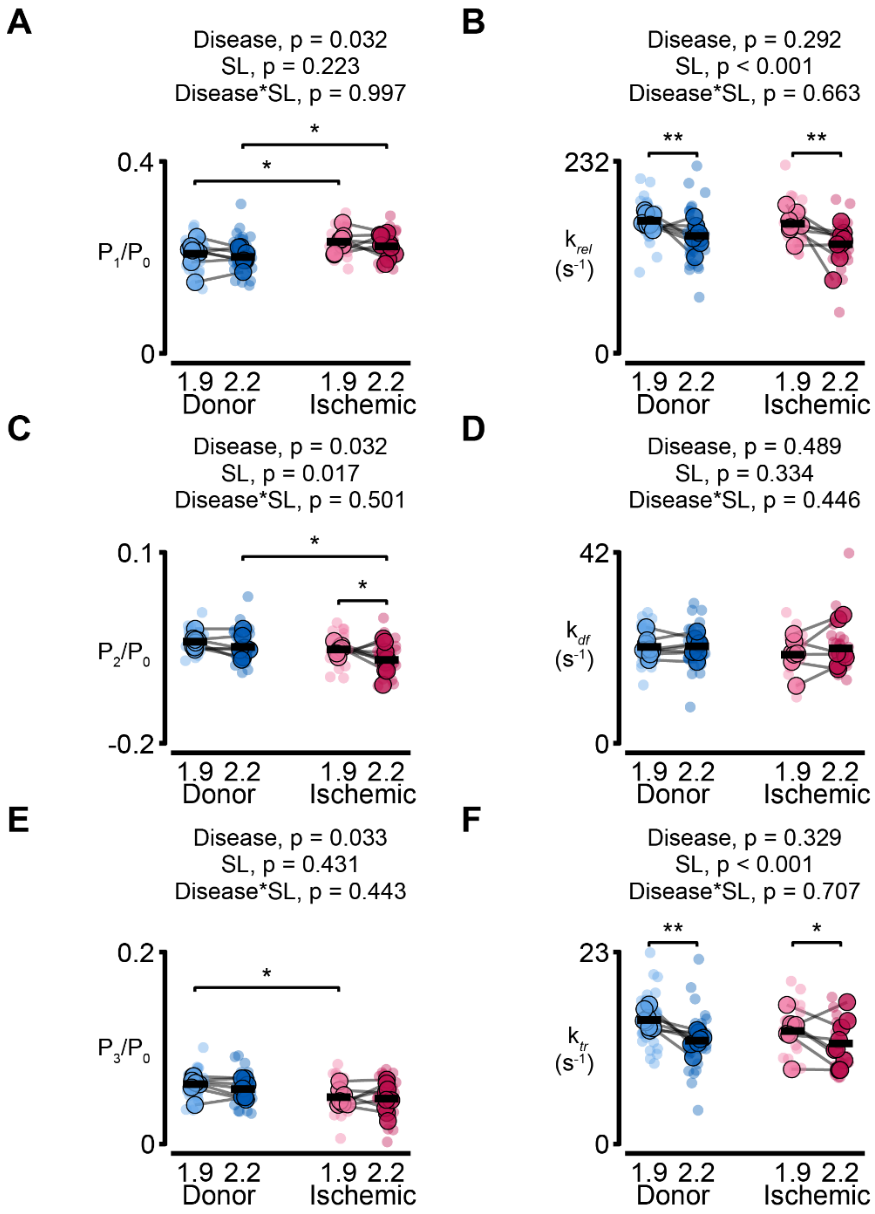


Supplementary Figure 3. Left ventricle cross-bridge kinetics are slower at longer sarcomere length

Parameters calculated from step-stretch force transients or ktr maneuver. Superplots show (A) P1, (C) P2, and (E) P3 normalized to P0 and (B) $k_{rel}$, (D) $k_{df}$, (F) $k_{tr}$. Data were analyzed using linear mixed models. *$p < 0.05$, **$p < 0.01$, ***$p<0.001$

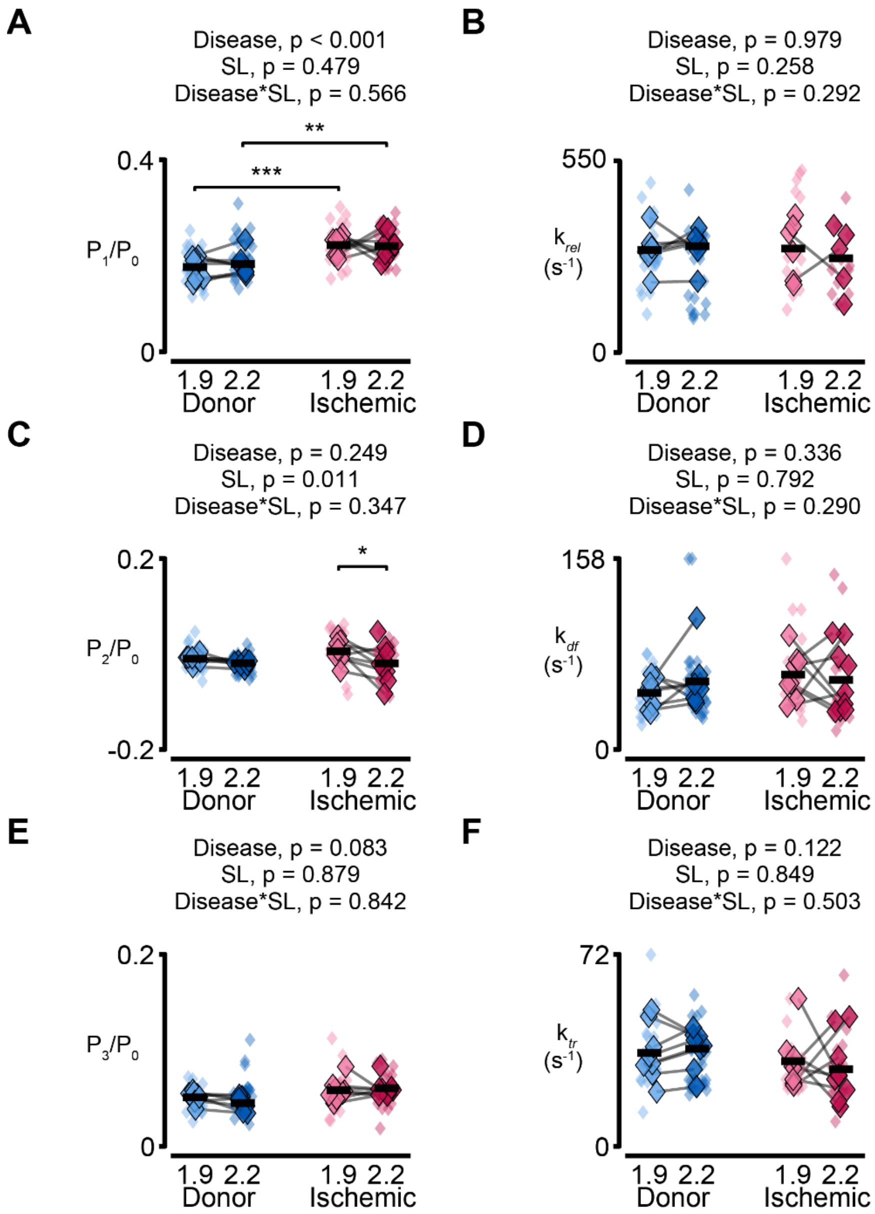


Supplementary Figure 4. Sarcomere length and disease status do not significantly alter left atrial cross-bridge kinetics

Parameters calculated from step-stretch force transients or ktr maneuver. Superplots show (A) P1, (C) P2, and (E) P3 normalized to P0 and (B) $k_{rel}$, (D) $k_{df}$, (F) $k_{tr}$. Data were analyzed using linear mixed models. *$p < 0.05$, **$p < 0.01$, ***$p<0.001$
